# Verification Debt in Automated Biological Research Workflows: A Proof-of-Concept

**DOI:** 10.64898/2026.09.29.755250

**Authors:** Shreya Gopalan, Sundaraparipurnan Narayanan

## Abstract

Recent biological research has shown significant interest and progress towards adoption of agentic AI systems. Domain specific agentic platforms with access to large collections of biological tools with code generation and/or querying capabilities, such as Biomni (illustrative case in point), are representative of the capability to orchestrate automated research workflows. Such platforms enable collaborative AI workspace for drug discovery, genomics, bioinformatics, literature research, and experimental planning for biologists and researchers. When such platforms are used for the aforementioned workflows, individually validating every agentic decision and the research process by which it was reached becomes challenging due to the scale of decisions involved and oftentimes, insufficient disclosure of assumptions and methodological choices, limited traceability of evidence, and interpretive judgments embedded throughout the workflow, giving rise to verification debt. Verification debt in agentic environments could result in unexpected failures and consequences in downstream experiments. This study focuses on empirical qualitative characterization of verification debt in agentic (prompt to outcomes) workflows. Biomni was chosen the testing AI agentic platform in this study mainly due to the descriptive traces it provides to track every agentic step. Verification debt is examined through two complementary angles: the individual agentic decision steps that carry claims and evidence, and the underlying research process by which the agent arrives at and reports them. This study attempts to characterize verification debt with 10 test cases (covering proteomics, genomics and drug discovery) with varying degree of biological nuance across three identical and independent runs. Verification debt is represented through 9 decision-step metrics and 13 research-process metrics, and also propose the VERIFY framework for evaluating scientific verifiability and validity in agentic workflows.

## 1 Introduction

Agentic AI environments are demonstrating growing capability for automating end-to-end biological research workflows. For instance, Virtual Lab introduced an end-to-end integration of scientific workflows with multiple LLM expert agents orchestrating each research component [1]. ToolUniverse, a massive undertaking integrating over 2,700 tools within an agentic ecosystem, democratized tool access and provided a platform for building customized AI scientists [2]. Biomni, a biomedical research agent, integrates more than 150 tools and 59 databases to execute complex, end-to-end scientific workflows, achieving human-level accuracy across multiple benchmarks [3].

Although such capabilities bring considerable potential for increasing the scale and speed of research, some studies have raised concerns that the magnitude and scale of automated research might push humans out of the loop [4, 5]. This reduction in human oversight could, in turn, increase verification debt. For the purposes of this paper, we define verification debt as the accumulation of agentic decisions and the research process by which it arrives at them, that a human researcher cannot fully verify due to the sheer volume and interconnected implications of the decisions themselves, or due to insufficient disclosure, evidentiary grounding, or traceability throughout the workflow [6, 7].

This study focuses on the qualitative empirical characterization of verification debt in agentic research platforms for biology. Biomni [3] was chosen as the agentic AI environment for this study, one of the reasons being the detailed trace records it provides within the user interface. This study evaluates verification debt from two angles: agentic decisions and the research process. Agentic decisions refer to the inferences, claims, and assumptions that the agent makes in the process of answering a research question. The research process refers to the methodological choices made by the agent, including the approach used to answer a research question, the choice of data/ database/ source, the inclusion and exclusion criteria, and the interpretation of analytical results.

## 2 Research Methodology

### 2.1 Experimental Setup

#### 2.1.1 Biomni Setup and Test Case Selection

The Biomni agentic environment includes a user interface through which users can submit research questions. With access to more than 150 standardized biological tools, 59 databases, and 105 software packages, the agent performs research tasks to answer these questions [3]. For this study, we formulated 10 high-level biological questions with varying degrees of biological nuance and requirements for validation, and instructed Biomni to run automated workflows to reach a conclusion for each question. The questions span protein function, structure, mutation, interaction, and druggability, as well as gene function, gene–disease association, transcriptomics, pathway analysis, and RNA–protein integration. Each question was run three times as identical and independent runs on Biomni to assess consistency, reproducibility, and fidelity. To ensure consistency in the experimental setup, we configured Biomni to use the default “Max” model. Follow-up clarification questions were answered automatically, and plan reviews were approved without changes. The research questions are described in Table 1.

**Table 1:** Research questions used to evaluate verification debt in Biomni and the rationale for their selection.

| No. | Domain | Candidate | Test question | Why this question |
| --- | --- | --- | --- | --- |
| 1 | Protein function | TP53 | What is the established biological function of TP53? | TP53 has extensive curated and experimental literature, making it a strong high-evidence baseline. |
| 2 | Protein structure | KRAS | How do KRAS structural domains relate to its function? | KRAS has well-characterized structures and functional domains, allowing verification across structural databases and literature. |
| 3 | Protein mutation | TP53 R175H | What are the functional consequences of TP53 R175H? | R175H has substantial functional literature but also requires distinguishing established evidence from mechanistic inference. |
| 4 | Protein interaction | BRCA1 | What are experimentally supported BRCA1 interaction partners? | BRCA1 has extensive interaction evidence, allowing comparison between curated, experimental, and predicted interactions. |
| 5 | Protein druggability | KRAS G12C | Is KRAS G12C a viable therapeutic target? | This question tests a long evidence chain from mutation → mechanism → druggability → clinical evidence; KRAS is also a well-established molecular testing target. |
| 6 | Gene function | PTEN | What is the biological role of PTEN? | PTEN has strong functional evidence but operates across multiple pathways, testing whether Biomni appropriately selects and weighs evidence. |
| 7 | Gene–disease association | APOE | What is the evidence linking APOE variants to Alzheimer’s disease? | This question requires Biomni to distinguish association, risk modification, mechanism, and causality, making it suitable for evaluating evidence and inference. |
| 8 | Transcriptomics | EGFR | Identify genes/pathways associated with EGFR-driven lung cancer. | EGFR provides a concrete disease context with established molecular biology and available cancer datasets, allowing assessment of dataset and method selection. |
| 9 | Pathway/network | TP53 | What pathways distinguish TP53-mutant from TP53-WT tumors? | TP53 provides a strong benchmark for testing enrichment-method choice and whether pathway association is incorrectly interpreted as mechanism. |
| 10 | RNA + protein | ERBB2/HER2 | Integrate RNA and protein data to determine whether HER2 is consistently altered in breast cancer. | HER2 is suitable because discordance between transcript and protein abundance is biologically meaningful and clinically relevant. |

#### 2.1.2 VERIFY Framework

This study was conceptualized around VERIFY, a framework that was developed to systematically characterize verification debt in agentic research pipelines. Verification debt was characterized from two angles: agentic decision steps and research process. Every metric used in this study assess one or more dimensions from the VERIFY framework.

##### V—Validity of the Research Question and Outcome

Assess whether the agent correctly interprets the user’s research question and translates it into an appropriate research objective or hypothesis.

##### E—Evidence and Epistemic Strength

Classify elements of the agentic research as information, claims, qualifiers, assumptions, interpretations, or hypotheses. For each element, assess the evidentiary basis, including direct observations, experimental or computational evidence, published literature, established methodological practice, agent inference, or assertion.

##### R—Research-Decision Appropriateness

Examine the appropriateness of the agent’s research decisions, including the selection of databases, tools, analytical methods, statistical approaches, parameters, thresholds, candidate-selection strategies, and experimental approaches. In the absence of appropriate candidates, mechanisms for recording uncertainty must be present.

##### I—Inference, Assumptions, and Scientific Integrity

Identify assumptions introduced during data analysis, interpretation, and hypothesis generation, and assess whether they are explicit, evidence-supported, and appropriate to the biological context.

##### F—Fidelity, Reproducibility, and Consistency

Assess whether the agent’s execution remains faithful to its stated intent, plan, or reasoning. Assess whether autonomous execution produces consistent research trajectories when the same research question is repeated. Where trajectories differ, assess whether the resulting approaches remain scientifically defensible and whether the differences introduce additional verification requirements.

##### Y—Yield and Verifiability of the Final Scientific Output

Assess the quality and adequacy of the final scientific output, whether it answers the original research question, whether its major conclusions are sufficiently supported by the preceding evidence chain, and whether the underlying claims, analyses, and decisions can be independently verified. Unresolved elements constitute the workflow’s verification debt, which an expert can validate individually.

### 2.2 Verification Methodology

Verification was performed at two levels: atomic-level agentic decisions and the research process.

#### 2.2.1 Agentic Decisions

An automated, LLM-driven extraction pipeline was developed to extract atomic-level agentic steps from the traces. Each atomic step or decision was classified as information, a claim, a qualifier, an assumption, an interpretation, or a hypothesis. The LLM-driven extraction was manually reviewed by one author to verify the extracted atomic steps and their corresponding classifications against the underlying execution traces and available biological evidence. Discrepancies identified during this review were examined and resolved through manual adjudication. The final dataset presented in the Supplementary data availability section (Supplementary A.1, Agentic-decision dataset) contains the human-verified and approved classifications only. Some well-known research questions (2 out of 10 questions) did not have traces at all. The extraction protocol broke down agent response into atomic-level steps in the absence of traces. This process ensured that the reported verification metrics reflected the underlying agentic workflow rather than the LLM verifier’s interpretation alone. Agentic decisions were assessed using 9 metrics identified by the authors to represent the feasibility and ease of verifying each atomic-level agentic step. Table 2 describes the metrics used in this study to assess atomic-level agentic decisions. The extracted atomic steps, classifications, and metric-level assessments constitute the *agentic-decision dataset*; access information is provided in Supplementary Section A.1.

**Table 2:**
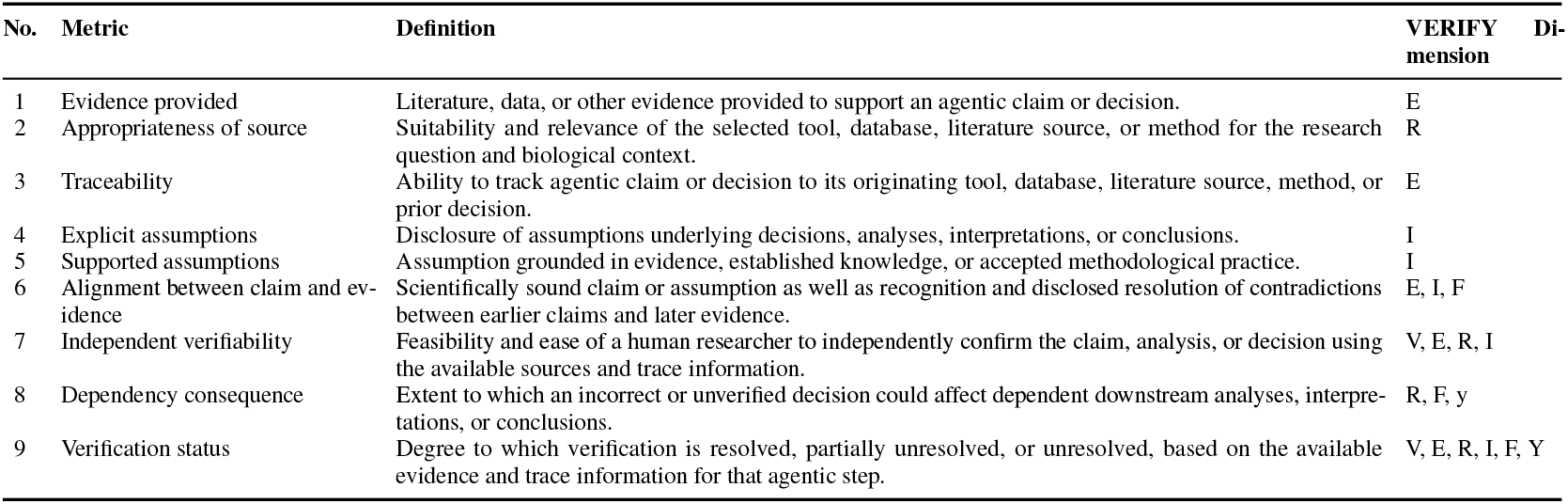
Definitions of the nine decision-step verification metrics used in this study, mapped to their corresponding VERIFY dimension(s)

For each of the 9 decision-step metrics, every atomic decision step across all three runs of a test case was scored on a three-point scale (Yes, No, Partial), assigned numeric values of 1, 0.5, and 0 respectively. Steps for which a given metric did not apply (e.g., assumption-related metrics on steps containing no assumption) were excluded from that metric’s denominator. The mean score across all applicable steps was then expressed as a percentage.

#### 2.2.2 Research Process

Within the scope of this study, the research process was reviewed using 13 metrics developed by the authors following an initial understanding of the results. Table 3 presents the metrics used to evaluate the research process adopted by the agent. The review deliberately excluded metrics that judged methodological choices that could reasonably depend on a researcher’s discretion when addressing a research question. For example, an agent might choose a literature search rather than a database such as KEGG for a question involving gene networks and pathways. However, insufficient disclosure of the limitations or disadvantages of such a choice was documented during review of the agent’s response.

**Table 3:**
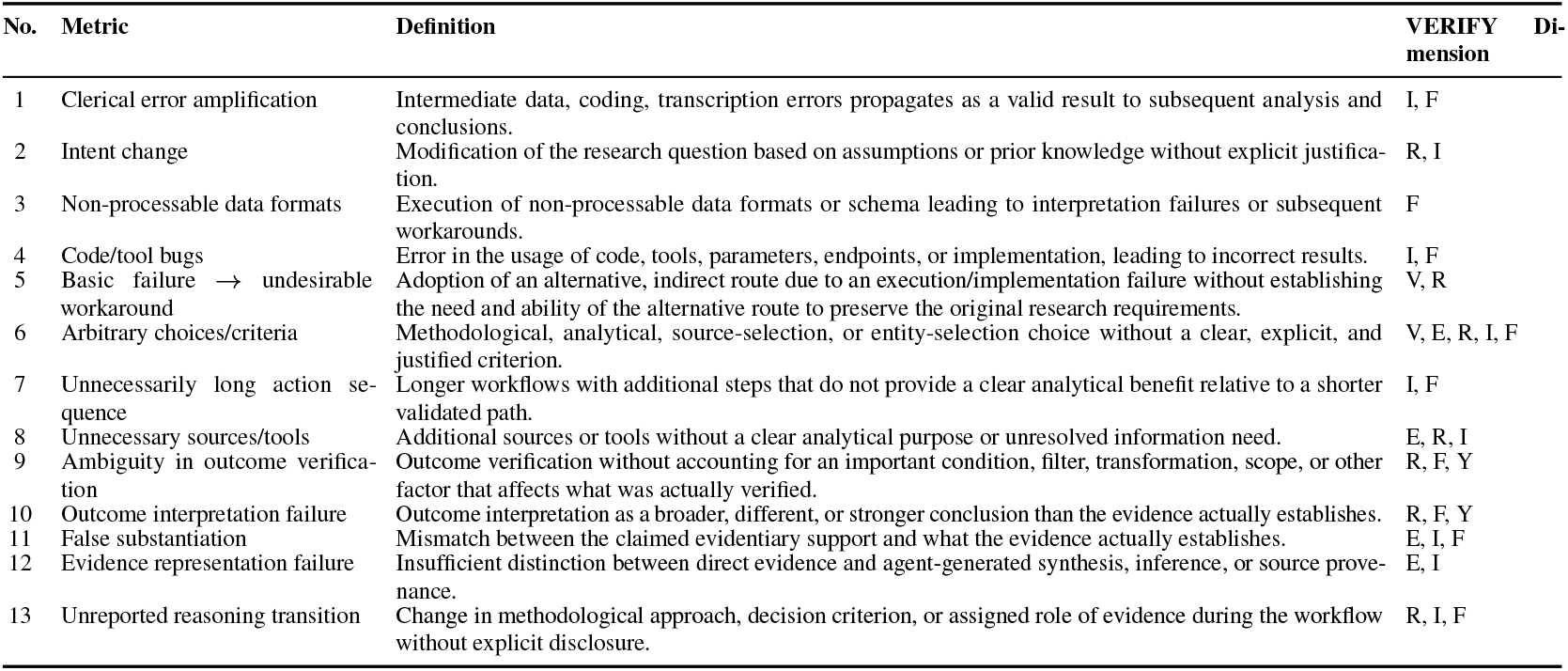
Definitions of the 13 research-process verification metrics used in this study, mapped to their corresponding VERIFY dimension(s)

Every test case was manually audited by one author to assess the research methodology adopted by the agent for each question. The manual review also considered how two or more interconnected trace steps contributed to a scientific conclusion. The agent’s traces and response report were reviewed to identify process-related inconsistencies that could lead to scientifically flawed or biased conclusions that a human researcher might be unable to verify without close supervision of each step and its downstream consequences. The resulting process-level audit records and assessments constitute the *research-process dataset*; access information is provided in Supplementary Section A.1.

Research process debt was recorded only where a methodological issue was identified in the trace, rather than scored for every step. For each test case, the reported percentage for a given issue type reflects the number of instances of that issue as a proportion of the total number of issues identified for that test case. Because this study’s issue identification was not exhaustive, these percentages should be interpreted as relative indicators of which issue types were most prevalent within each test case’s identified issues, rather than as absolute rates across all steps performed.

## 3 Results and Discussion

Across the 10 questions, we assessed verification debt from two complementary angles: debt arising at the level of individual agentic decision steps, and debt arising from the agent’s research methodology. As question complexity increased, both the number of decision steps and the number of process-related inconsistencies increased correspondingly. While this could imply that decision steps are directly proportional to the number of process inconsistencies, since the sample size was too small and this analysis is not exhaustive, it cannot be confirmed if there is a correlation between these two dimensions.

The research approach across all three runs for a question predominantly remained the same, with very few exceptions. However, research rigor and depth appeared to vary between the three runs. For example, the differences we observed were in terms of number of literature, choice of databases, exploratory vs targeted analysis, number of steps for an identical question, interpretation of the question has slight modulations across runs, choice to rely on prior knowledge Vs. literature, and its confidence, presentation of answers with or without caveats and references. Given the dynamic nature of agent runs and inability to attribute root causes for such variations, this study is not quantifying the implications of such variance from a verification debt perspective.

Across the 10 questions, 53% of the agentic decisions or steps were on claims, 8.6% were interpretations of results or information, and 7.6% consisted of scientific information from various sources. The rest of them were assumptions made by the agent. Notably, the majority of these claims were cited and the sources were typically scientific studies (with one exception), leading to majority of the verification debt in agentic decisions to be concentrated in explicit disclosure of assumptions and dependency consequence.

For some runs, especially for research questions that were well-known, there were no traces or even cited references. This implies that the agent tends to answer well-known biological questions from its own prior knowledge without explicit disclosure that it is doing so, leaving users with no way to verify the source of information. This occurred in 2 of the 10 questions. Also, in some cases the filter criteria used for literature searches varied (e.g. number of relevant papers selected and search period) across runs, with no reasoning in the traces as to why such variation existed. This inconsistency was observed across almost all of the questions, making it one of the more systemic sources of process debt identified in this study.

### 3.1 Metric-Level Analysis of Agentic Decision Debt

Aggregate-level trends are mentioned below across the 9 metrics to quantify verification debt in agentic decision steps. The heatmap (Figure 1) shows a breakdown of these metrics across the test cases (research questions).

**Figure 1.**
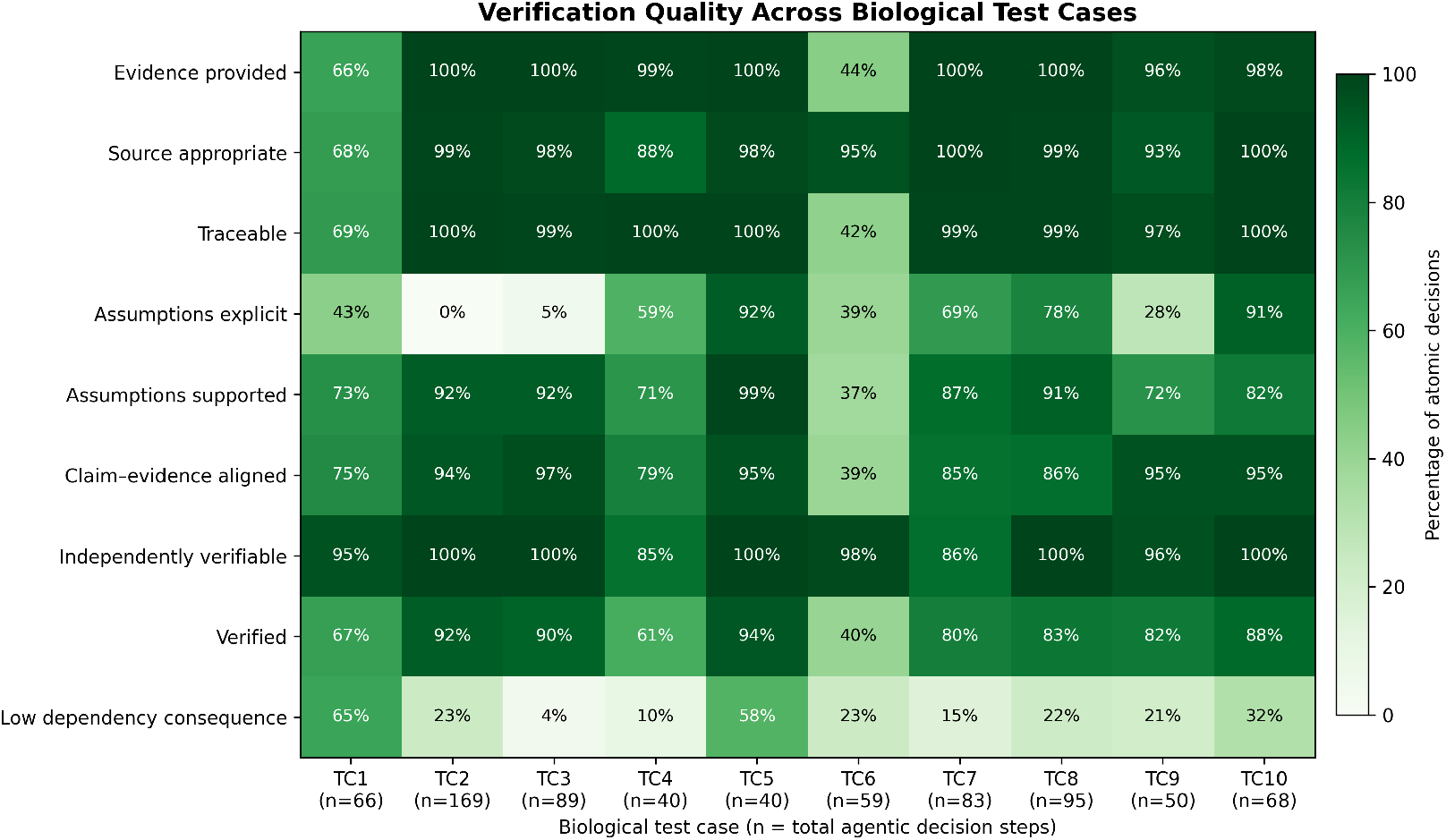
Verification quality across the agentic-decision metrics and biological test cases. Mean verification score per metric (0–100%), averaged across all atomic decision steps for each test case (across 3 runs). N/A steps to which a metric did not apply were excluded from that metric’s denominator.

Verification debt related to evidence availability, appropriateness of source, traceability of evidence was low for most questions, with 8 of 10 test cases scoring 88% or higher on all three metrics. The only outliers were Test case 1 (TC1) and TC6, which were well-known biological questions, and therefore citations and traces were absent in some of the runs. This implied that, for the majority of test cases, the agent had appropriate evidences that could be traced back to the source.

Assumptions handling showed a broader spread of verification debt across test cases. Explicit disclosure of assumptions ranged from as low as 0% (TC2) and 5% (TC3) to over 90% (TC5, TC10), indicating that whether an agent disclosed its assumptions as assumptions, rather than presenting them as cited claims varied substantially across test cases. Assumption support was comparatively stronger and more consistent, with most questions at 70% or above, though TC6 again lagged at 37%. The openly-declared assumptions, were however, supported with scientific grounding than random assumptions without data/evidence-backing. Most test cases showed a higher degree of supported assumptions (over 70%), with only TC6 lagging at 37% probably due to the lack of any trace or citation. Inconsistencies in handling of assumptions showed a trend where if the agent classified some step as an assumption, it was mostly supported through prior steps or well-known science. However, if the agent did not realize/disclose it was making an assumption, there are chances that the assumption was not strongly supported.

Alignment of claims with evidence (literature or database) was generally strong, ranging from 75–97% for 9 of 10 questions, with TC6 as the sole exception at 39%. This could indicate that majority of the agentic claims were based on prior knowledge from training, where the agent knew what to search for and where, rather than fresh research. Independent verifiability showed a solid trend with every question at 85% or above, correlating strongly with evidence availability and traceability since every claim can be independently verified, if provided with a source/citation The most consistent verification debt across the entire set of test cases were concentrated in dependency consequence. Unlike every other metric, where most questions clustered at the low-debt end, dependency consequence showed the opposite pattern with 8 of 10 questions had 32% or fewer decisions with low, none, or local downstream consequence, implying that the majority of decisions in nearly every question carried a critical, high, or propagating dependency risk. Only TC1 (65%) and TC5 (58%) showed a majority of decisions with comparatively lower consequences. Dependency consequence is the only metric across the 10 metrics that measures the consequence of lack of verification, implying that almost every agentic step has a downstream propagating effect, if left unchecked.

Verification status, for the given set of 10 test cases, was generally strong scoring 80 - 94% in 7 out of 10 test cases. TC1 and TC4 had a low verification status (67% and 40% respectively) due to the lack of traces and evidence, leading to reconstruction of provenance by the human reviewer for verification. TC4 also has a low verification status (61%) due to methodological inconsistencies and subsequent agentic decisions that could not be validated. Verification status is a human-reviewer-assessment metric capturing the overall degree to which each agentic decision step, across all test cases, can be considered verified, which is informed by but not strictly derived from the other 8 metrics. For this dataset of 10 test cases, the human review found majority of the agentic decision steps verifiable.

### 3.2 Metric-Level Analysis of Process Debt

Across the 10 test cases, 105 instances of process-oriented verification debt were identified, spanning 13 metrics. Figure 2 shows the breakdown of these metrics across the 10 test cases. The number of issues varied substantially, 3 in TC3 to 24 in TC10, displaying a correlation between the complexity of the test case and number of identified issues. This analysis is not exhaustive, and therefore, the identified methodological issues might not represent the actual spread of process inconsistencies found. Therefore, the percentages mentioned here must not be considered absolute, instead it should be considered relative, providing an idea of the type of process-oriented verification debt that are most commonly found.

**Figure 2.**
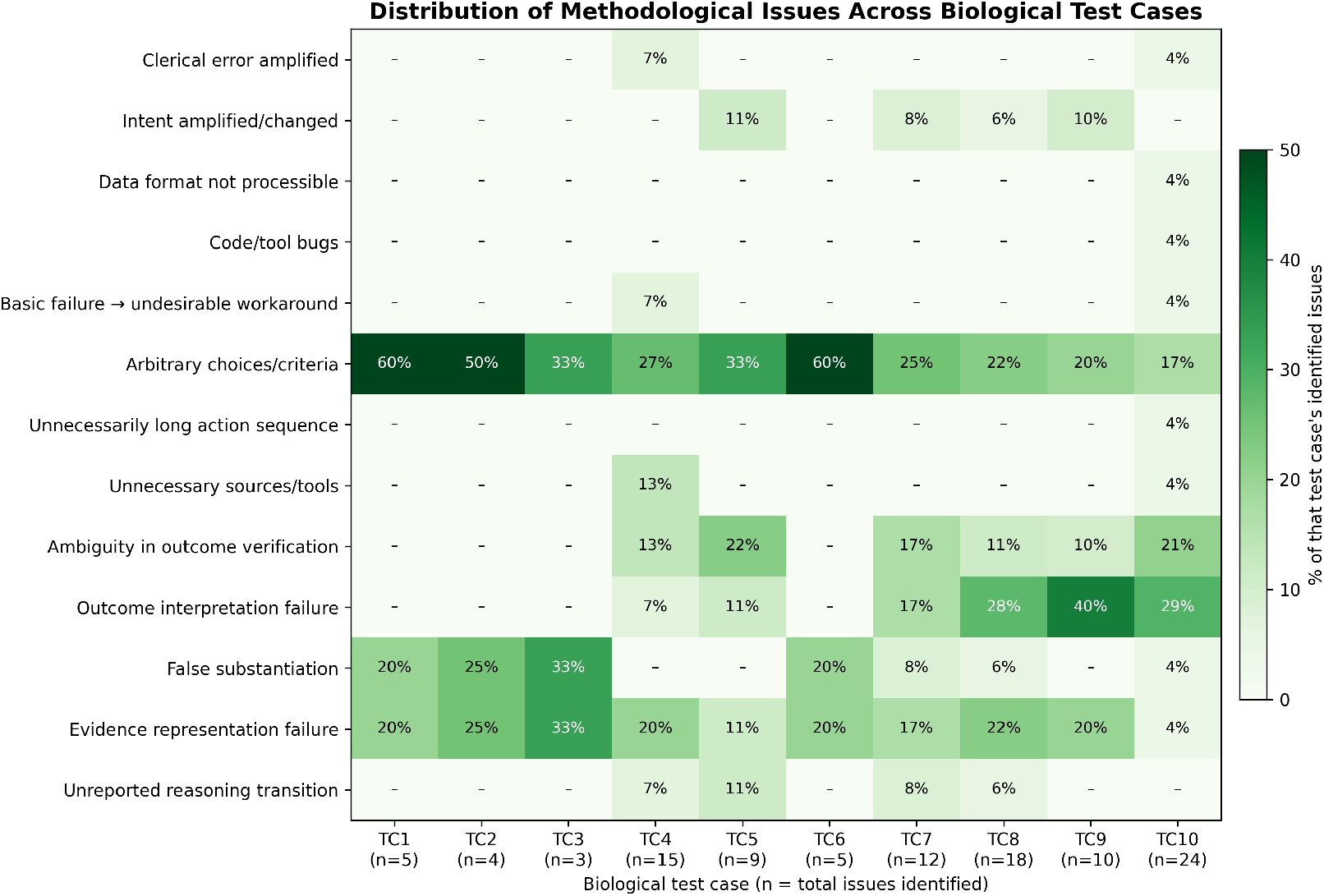
Verification quality across research process metrics and biological test cases. Proportion of each test case’s identified methodological issues attributable to each of the 13 issue types; percentages are relative to that test case’s total identified issues.

The most common process debt across the test cases was Arbitrary choices/criteria. TC1 and TC6 had the highest percentage of debt due to arbitrary choices/criteria (3 of 5 issues in each case), and was found to be prominent even in the two largest test cases (TC9: 20%, 2 of 10 and TC10: 17%, 4 of 24). This issue indicated that in majority of the test cases, methodological choices were not explicitly disclosed including ranking mechanisms of results, filtering criteria, source selection, and use of training data without explicit mention, to name a few.

False substantiation occurred in all the well-known test cases (TC1, TC2, TC3, and TC6), where the cited literature did not actually contain the information. This was because the agent typically receives triage one-line summaries of a publication, and in well-known cases, it tended to cite some literature that looks valid from the summaries, without actually checking if the citation contains the relevant content. This was generally the case for any literature that was cited by the agent, where at the beginning, it receives a triage summary, and if it finds the summary relevant and the question adequately complex, the agent decides to read the abstract. This becomes concerning because there is no disclosure from the agent to the level of information it has accessed in the agent response/report or the traces, leading to evidence representation failures.

Evidence representation failure is a debt that occurred consistently but manifests differently for different test cases. For the simpler, and well-known test cases (TC1-TC3, TC6), it manifests as an issue relating to lack of explicit disclosure to information that the agent has accessed. For the complex test cases, evidence representation failure manifests as granularity mismatch in citations (citations consider the contents of the whole paragraph than to the specific claim that they are cited for), evidence strength conflation (mouse studies and GWAS presented in the same claim without distinction between their evidence strength), evidence reconciliation issues (Agent response contains citations, but information from the citations are not used anywhere in the analysis) and omission/underrepresentation (some analysis angles were pursued and dropped/less represented without any explicit reasoning).

Outcome interpretation failures occurred mostly in complex and high decision step test cases, with 28 to 40% of issues in TC8, TC9, and TC10 (5 out of 18, 4 out of 10, and 7 out of 24 respectively). Ambiguity in outcome verification followed a similar trend, with 10 to 22% of issues in high complexity cases. These metrics suggest that as task complexity increased, verification debt shifted from disclosure-related failures towards interpretative failures, with the agent drawing broader or stronger conclusions than the evidence supported or failing to verify what a result actually established.

The remaining 7 metrics (clerical error amplification, intent change, unprocessable data formats, code/tool bugs, undesirable workaround failures, unnecessarily long action sequences, and unreported reasoning transitions) occurred rarely (1–4 instances each across the full dataset) and were concentrated almost entirely in the complex cases, again iterating the fact the debt shift to interpretative failures in complex cases.

### 3.3 Discussion

Overall, our findings indicate that Biomni, as a biomedical research agent, is capable of conducting rigorous research workflows. The agent could retrieve literature evidence, select appropriate databases, provide reasoning traces, track and correct some of its own errors, and generally produce claims that align with the available evidence. Traces were also, for the majority, accessible enough for a domain specialist to review and audit, which is a necessity for any form of external verification.

However, the findings also highlight certain important caveats, at both the decision-step and process levels. At the level of agentic decision steps

- Literature retrieval often relied on a search API returning only a summary of an abstract, rather than the complete abstract or full text, meaning claims could be grounded in less information than they appeared to be.
- Reasoning traces were, in some cases, incomplete or unavailable for individual decision steps, creating difficulties in tracing agentic steps.
- Assumptions, including both evidence-supported reasoning and more speculative inferences, may not always be explicitly disclosed.
- Literature grounding may rely on non-standard or non-primary sources.
- Scientific values and statistical analyses may not always be derived using standard procedures. In some instances, analytical thresholds or methods appear to have been selected retrospectively based on the observed data distribution. Crucial supporting information, including effect sizes and confidence intervals, may also be omitted. This finding was also reflected in [8] where the authors demonstrate two agents with opposing goals produce statistically defensible but contradictory conclusions with the same dataset through differing analytical choices.

At the level of research process,

- The agentic process, even when traces were available, is largely opaque in terms of particular methodological path was selected over alternatives.
- Agentic responses also contained numerous loose ends in how evidence was represented, what was disclosed, and what was performed but never explicitly mentioned.
- Finally, the agent did not always verify every aspect of an outcome before presenting it as final, and its interpretation of outcomes was at times flawed, leading to the interpretative failures discussed above.

One of the most important findings of this study is that these issues do not occur consistently. Rather, they emerge unpredictably across different points of the research pipeline, and even across different runs of the same research question, making them difficult to anticipate or systematically track. This unpredictability is further compounded by the fact that different research questions require different degrees of domain-specific judgment and contextual knowledge. Determining whether a given claim is scientifically valid for a particular scenario may itself require expert manual review, rather than a generic verification protocol.

These findings are also broadly consistent with studies evaluating agentic AI for biological research. DrugDiscoveryBench [9], which evaluates agents on early-drug-discovery tasks, found that the dominant failure mode across frontier models was not retrieval or tool access but domain-specific reasoning (agents applying an incorrect scientific premise or misinterpreting available data even when their inputs and tools were correct), accounting for over 45–61% of failures across models. Similarly, BioAgent Bench (BioAgent Bench: An AI Agent Evaluation Suite for Bioinformatics) [10] demonstrated that high pipeline completion rates coexisted with brittle step-level reasoning under input perturbations and high run-to-run output variability across trials. These limitations were further observed in single-cell multi-omics using the M3A Framework [11], where autonomous agents performed well on canonical cell types but failed on rare cell populations (50.8% cross-family errors) and exhibited thematic literature bias by over-ranking heavily documented cell programs over under-documented novel drivers.

This study was designed as a proof of concept to establish the presence of verification debt in automated research pipelines in biology and has certain limitations. The sample of 10 research questions is too small to quantify verification debt comprehensively or support a rigorous assessment of its downstream impact. An LLM was used to decompose workflows into atomic decision steps and identify instances of the evaluation metrics, followed by manual validation of the resulting annotations. Nevertheless, because the initial analysis was LLM-assisted rather than based on exhaustive manual review, some context-dependent or domain-specific issues requiring expert judgment may have been missed. Additionally, some code-related issues might not have been identified in this study, since the scope was to determine verifiability of the biological intent and research direction rather than computational. Similarly, this study also did not attempt query modifications to see if agent responses changed with additional context.

## 4 Conclusion

Agentic AI systems and automated workflows are being adopted increasingly in biological research. This study identifies inconsistencies in agentic research processes that warrant expert intervention at multiple checkpoints across the research pipeline. Although the speed and scale of these systems offer substantial potential to accelerate research, they also create a growing gap between automated execution and the capacity for human review and validation.

This study attempts to characterize verification debt in agentic biological research pipelines, examined through two complementary angles: agentic decision steps claim and research process. Across both angles, our findings show that verification debt does not accumulate uniformly or predictably. It is concentrated in different metrics for different questions, shifting from disclosure-type gaps in simpler, well-known questions to interpretative failures in more complex ones. Verification debt cannot be reliably anticipated from a every individual step alone or by evaluating outputs or processes in isolation from one another.

These findings motivate the case for a systematic verification protocol that evaluates agentic research pipelines along both angles examined in this study - the individual decision steps that carry claims, evidence, and assumptions, and the research processes that determines how those claims are arrived at and represented. A pipeline may produce individually well-cited claims while the process that generated them remains opaque, arbitrary, or inconsistent across runs, or, conversely, may follow a defensible methodological process while individual claims still fail to hold under independent scrutiny. A complete account of an agentic system’s reliability requires both.

Ultimately, this study underscores that transparency (presence of traces, citations, and disclosed reasoning) is a necessary condition for verification, but not a sufficient one. An agentic pipeline can be fully traceable and still produce claims, assumptions, and methodological choices that a domain expert, upon review, would find unreliable or insufficiently supported. This study emphasizes that verification debt cannot be resolved by transparency alone. It requires deliberate, structured evaluation of what is being claimed, how it was reached, and whether it can withstand independent scrutiny.

## 5 Future directions

This study characterizes verification debt through a framework, VERIFY. Future research must build on frameworks like VERIFY to move from characterization to quantification, establishing measurable, operational protocols for each dimension such that verification debt can be systematically evaluated, measured, and scored across agentic systems rather than only characterized and described. We propose VERIFY as a starting point for this transition: a structural foundation for future work seeking to operationalize verification debt assessment as a standard, quantifiable component of agentic research pipeline evaluation, applicable beyond the biological domain examined here. This study explores the VERIFY framework for both angles: agentic decision-step and research process through majorly manual validation. The manual review process used in the present study is not scalable, and any automated or semi-automated alternative (an LLM-based judge) would itself introduce a new verification question on whether the judge’s own assessments can be trusted. Therefore, future studies should also focus on systematic scalable procedures to verify large scale research pipelines.

## A Supplementary Material

### A.1 Data Availability

The two datasets supporting this study are reported separately to reflect the two levels of verification described in the research methodology:

#### Agentic-decision dataset

Atomic agentic steps, decision classifications, and assessments across the 10 verification metrics are available on Figshare at https://figshare.com/s/ca12026c9f6ed046d352.

#### Research-process dataset

Process-level audit records and assessments across the 13 research-process metrics are available on Figshare at https://figshare.com/s/ab2b33817aab0bf9ca3.

## References

[1] Kyle Swanson, Wesley Wu, Nash L. Bulaong, John E. Pak, and James Zou. The virtual lab of AI agents designs new SARS-CoV-2 nanobodies. Nature, 646:716–723, 2025. doi: 10.1038/s41586-025-09442-9. URL https://doi.org/10.1038/s41586-025-09442-9.

[2] Shanghua Gao, Richard Zhu, Pengwei Sui, Zhenglun Kong, Sufian Aldogom, Yepeng Huang, Ayush Noori, Reza Shamji, Krishna Parvataneni, Theodoros Tsiligkaridis, and Marinka Zitnik. ToolUniverse: An open platform for democratizing AI scientists, 2026. URL https://arxiv.org/abs/2509.23426.

[3] Kexin Huang, Serena Zhang, Hanchen Wang, Yuanhao Qu, Yingzhou Lu, Ryan Li, Yusuf Roohani, Lin Qiu, Shiyi Cao, Gavin Li, Junze Zhang, D. Yin Rick Wierenga, Deniz Kavi, Sherry Liu, Tianwei She, Shruti Marwaha, Jennefer N. Carter, Xin Zhou, Matthew T. Wheeler, Jonathan A. Bernstein, Mengdi Wang, Peng He, Jingtian Zhou, Michael P. Snyder, Le Cong, Aviv Regev, and Jure Leskovec. Autonomous biomedical research with an artificial intelligence agent. Science, 393(6813):eadz4351, 2026. doi: 10.1126/science.adz4351. URL https://www.science.org/doi/10.1126/science.adz4351.

[4] Belinda Mo. The age of AI agents demands a new scientific paradigm to sustain trustworthy science, 2026. URL https://arxiv.org/abs/2607.26064.

[5] Cristina Cornelio, Takuya Ito, Ryan Cory-Wright, Sanjeeb Dash, and Lior Horesh. The need for verification in AI-driven scientific discovery, 2025. URL https://arxiv.org/abs/2509.01398v2.

[6] Yury Korolev. The year of silent failures: AI technical debt and the crisis of verification. AI Decisions, 1(2):5–22, December 2025. doi: 10.65114/aide.gmry6g41. URL https://doi.org/10.65114/aide.gmry6g41.

[7] Christian Catalini, Xiang Hui, and Jane Wu. Some simple economics of AGI, 2026. URL https://arxiv.org/abs/2602.20946.

[8] Dionizije Fa and Marko Culjak. Sound agentic science requires adversarial experiments. In ICLR 2026 Workshop on Agents in the Wild, 2026. doi: 10.48550/arXiv.2604.22080. URL https://arxiv.org/abs/2604.22080.

[9] Afra Feyza Akyürek, Xinming Tu, Alec Gutmanstein, Jason Qin, Divyansh Agarwal, Sofia Monasdotter, Sergey Chekhov, Brenda Hernandez Villegas, Kirill Chugunov, Judah Engel, Veronica Chatrath, Oscar Kavanagh, Geobio Boo, Ernesto Hernandez, Ying Liu, Emily Xue, Aakash Sabharwal, Daniel Yue Zhang, Zainab Doctor, Yuanhao Qu, Yunzhong He, and Sami Hassaan. DrugDiscoveryBench: Can coding agents assist early-stage drug discovery? Scale Labs, June 2026. URL https://static.remotasks.com/uploads/67f465178e967e5b6889c886/DrugDiscoveryBench.pdf. Accessed: 2026-09-25.

[10] Dionizije Fa, Marko Čuljak, Bruno PandŽa, and Mateo Čupić. BioAgent Bench: An AI agent evaluation suite for bioinformatics, 2026. URL https://arxiv.org/abs/2601.21800.

[11] Shreya Johri, Erica Pimenta, Josephine Yates, Jingxin Fu, Erik L. Bao, Hyeji Jun, Brendan Reardon, Sasha Bacot, Maha Shady, Doris Fu, Wenbin Mei, Sabrina Y. Camp, Jihye Park, and Eliezer Van Allen. Evaluating agentic AI for biological discovery in autonomous and copilot settings. bioRxiv, June 2026. doi: 10.64898/2026.06.04.729919. URL https://doi.org/10.64898/2026.06.04.729919.

